# Integrative Proteomic Analysis of Posttranslational Modification in The Inflammatory Response

**DOI:** 10.1101/2020.07.20.212134

**Authors:** Feiyang Ji, Menghao Zhou, Huihui Zhu, Zhengyi Jiang, Qirui Li, Xiaoxi Ouyang, Yiming Lv, Sainan Zhang, Tian Wu, Lanjuan Li

## Abstract

The posttranslational modification (PTM) of proteins, particularly acetylation, phosphorylation and ubiquitination, plays a critical role in the host innate immune response. PTMs’ dynamic changes and the crosstalk among them are complicated. To build a comprehensive dynamic network of inflammation related proteins, we integrated data from the whole cell proteome (WCP), acetylome, phosphoproteome and ubiquitinome of human and mouse macrophages. Our datasets of acetylation, phosphorylation and ubiquitination sites helped identify PTM crosstalk within and across proteins involved in the inflammatory response. Stimulation of macrophages by lipopolysaccharide (LPS) resulted in both degradative and non-degradative ubiquitination. Moreover, this study contributes to the interpretation of the roles of known inflammatory molecules and the discovery of novel inflammatory proteins.

## Introduction

Macrophages are resident phagocytic cells which act as effector cells in innate immune system and play a crucial role in initiating adaptive immunity by means of recruiting other immune cells [1]. Located throughout the tissues in the body, macrophages are among the first defensive cells to interact with foreign or abnormal host cells and their products. They are the most efficient phagocytes that can ingest and process foreign materials, dead cells and debris; they also release various secretory products to mobilize other host cells and influence other resident cells in the inflammatory response [2]. Posttranslational modifications (PTMs) have been reported to play a critical role in regulating multiple inflammatory signalling pathways. Phosphorylation, polyubiquitination, and acetylation exert a diversity of effects on pathogen recognition receptor(PRR)-dependent inflammatory responses [3].

Phosphorylation is a widely investigated type of PTM in innate immunity [4]. It is catalysed by protein kinases and reversed by protein phosphatases. The phosphorylation and dephosphorylation of certain proteins regulate the activation and deactivation of many TLR-dependent signalling molecules. Typical examples are *MAPKs, IκBα, IKKα, IKKβ*, and *IRF3* [5]. Existing research also recognizes the critical role of the phosphorylation of innate immune adaptor proteins. For example, the phosphorylation of *MAVS, STING*, and *TRIF* is necessary for *IRF3*’s recruitment and type I *IFN*’s production, which is essential for the activation of antiviral immunity [6]. Protein ubiquitination also plays a pivotal role in a wide variety of immunological processes [7]. Ubiquitin, a highly conserved polypeptide which comprises 76 amino acids, is attached to substrates by a complicated three-step enzymatic cascade [8]. Seven lysine residues within ubiquitin can be bonded with poly-ubiquitin chains, namely, K6, K11, K27, K29, K33, K48, K63. Ubiquitin can also be ubiquitinated at the N-terminal methionine residue (M1) [3, 9]. Different types of ubiquitin connections lead to different outcomes. For example, K48-linked polyubiquitination of *IκB* results in its proteasomal degradation, promoting the nuclear translocation of *NF-κB* [10]. In contrast, K63-linked polyubiquitination of *TAB2/3, TRAF6, NEMO*, and *TRAF3* are “proteasome-independent” and requisite for activating *NF-κB* and *IRF3* [3]. Furthermore, different linkage types of ubiquitin chains, such as M1, K11-, K48-, and K63-linked ubiquitin chains, are both involved in the TNF-induced inflammatory signalling pathway and play critical parts in the regulation of the downstream signalling cascade [8]. In addition to those two widespread PTMs, several studies have highlighted important roles for other PTMs, including acetylation, in the regulation of immune responses. The change of chromatin structure via the acetylation of histones and its influence on gene transcription are well understood [11, 12]. For example, in antiviral immunity, *HDAC9* is upregulated and in turn deacetylates the kinase *TBK1* for activation, leading to increased *IFN* production [13]. Meanwhile, acetylation has been discovered in many non-histone substrates, which participate in immune system [14].

In the past few years, evidences for comprehensive crosstalk between PTMs has accumulated. An example of this crosstalk is the *NF-κB* signalling pathway, where *IKKβ* phosphorylates *IκBα*, resulting in the K48-linked ubiquitination and degradation of *IκBα*, the subsequent release of *NF-κB* and the entry of *p65* and *p50* dimers into the nucleus to activate target genes. As a matter of fact, some E3 ubiquitin ligases must be phosphorylated to become catalytically active or only ubiquitylate their substrates when the latter are phosphorylated [15]. According to a recent study by Huai et al, all three PTMs mentioned above occur in the same protein. In more detail, interferon regulatory factor 3 (*IRF3*) activity is strongly regulated by PTMs, such as ubiquitination and phosphorylation; however, *KAT8* inhibits antiviral immunity by acetylating *IRF3* [16]. The combination of various PTMs on protein apparently creates a “PTM code” that is recognized by specific effectors to initiate or inhibit downstream events [17].

Although various studies have investigated the role of PTMs in innate immunity, few quantitative analyses of PTMs have been conducted in this area. Furthermore, the crosstalk between different PTMs is always a difficult subject to explore, as the PTMs usually occur too rapidly to measure. This study seeks to obtain data that will help address these research gaps. We selected two types of macrophages, Raw macrophages from mouse and Thp1 macrophages from human, to establish the model of the inflammatory response induced by LPS stimulation. We applied high-resolution mass spectrometry to discover and quantify the changes in the three types of PTMs during innate immune responses at different time points. Using the advanced stable isotope labelling with amino acids in cell culture (SILAC) technique, we successfully profiled all three modifications with high accuracy and precision [18]. We measured the changes in PTMs at 30 minutes and 2 hours after LPS stimulation, and then used the unsupervised clustering method to divide PTM events into different groups for subsequent analysis. The experimental analysis presented here represents one of the first investigations into PTM crosstalk during the innate immune response. Our findings could make an irreplaceable contribution to the field of novel changes in PTMs occurring during inflammation.

## Results

### The workflow of the integrative proteomics analysis of LPS-stimulated macrophages

The transmission of signals from inflammatory signalling pathways involves the PTM of various proteins. We examined the changes in the acetylome, phosphoproteome and ubiquitinome in two types of inflammatory cells (Raw and Thp1) at different time points after LPS stimulation to investigate the relationship between the PTMs of various proteins during this process. We performed quantitative profiling of acetylation sites using Ac-Lys proteomics. Ac-Lys peptides result from tryptic cleavage of an acetylated lysine substrate and can be enriched with an Ac-Lys antibody. TiO_2_, Fe-NTA and P-Tyr methods were used to quantify the changes in the phosphoproteome. TiO_2_ and Fe-NTA are complementary methods to enrich phosphorylation sites in serine, threonine and tyrosine substrates. In contrast, a P-Tyr antibody specifically enriches phosphorylation sites in tyrosine substrates. We also performed quantitative profiling of ubiquitination sites using K-ε-GG proteomics. K-ε-GG remnants are a result of tryptic cleavage of a ubiquitinated lysine substrate and are enriched by a K-ε-GG antibody. We applied SILAC for the relative quantification of PTM sites, and all experiments were performed in three biological replicates. In addition, we increased the number of identified proteins and PTM sites using fractionation methods (**Figure 1**). After comparing the three phosphorylation omics methods, approximately one-third to half of the phosphorylation sites were quantified using both TiO_2_ and Fe-NTA methods. Meanwhile, most of the phosphorylation sites quantified using P-Tyr method were different from the sites identified with the other two methods (Figure S1A). The effects of MG132 on the identification of protein and ubiquitination sites were compared, and MG132 had little effect on the identification of proteins in the whole cell proteome (WCP), but exerted a relatively significant effect on the identification of ubiquitination sites in the ubiquitinome (Figure S1B).

**Figure 1.**
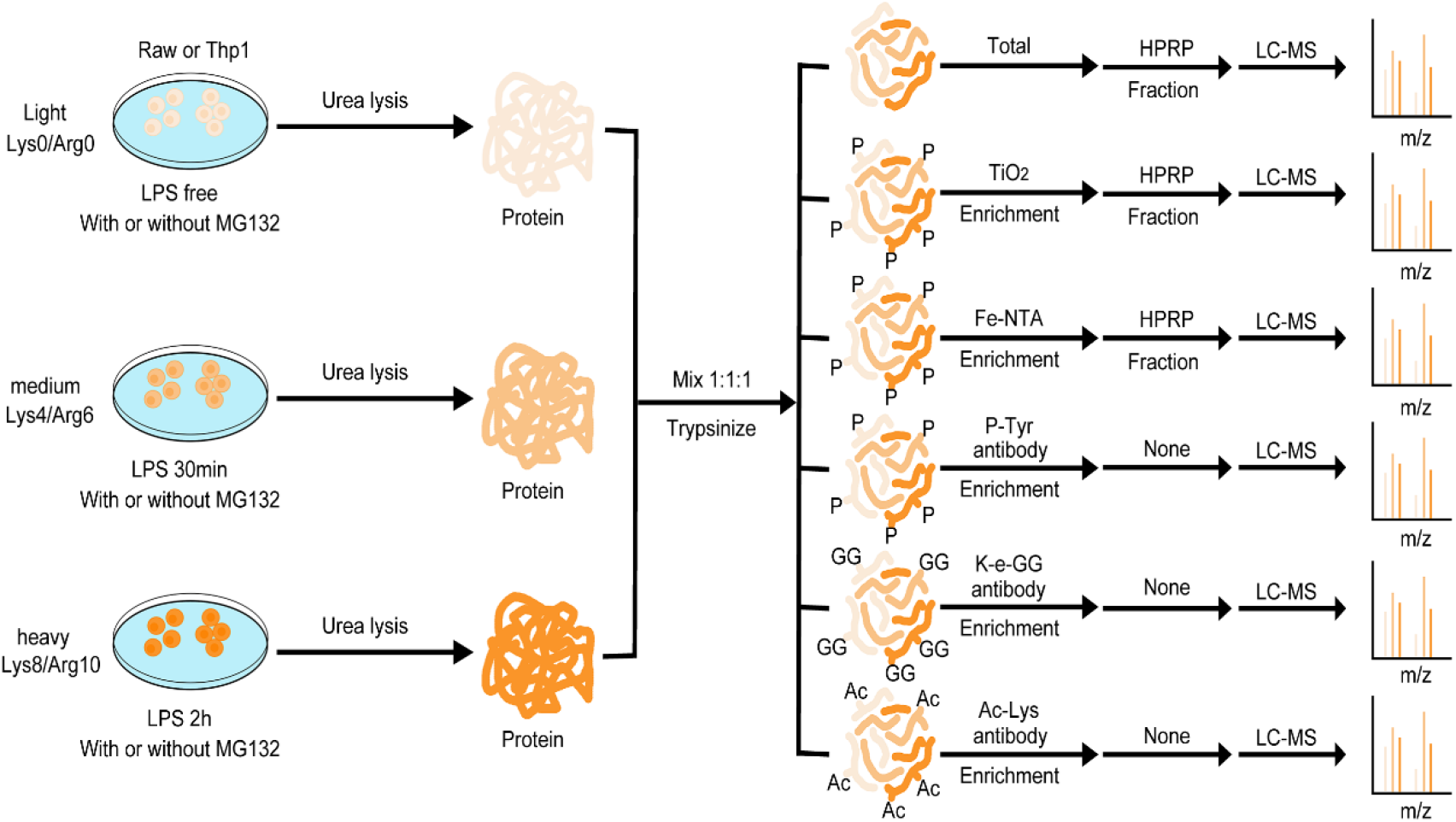
Integrative proteomics analysis of macrophages stimulated with LPS. Schematic of the integrated proteomics workflow: Raw and Thp1 cells cultivated in light, medium or heavy “SILAC, stable isotope labelling with amino acids in cell culture” medium was stimulated with “LPS, lipopolysaccharide” at different time points the presence or absence of MG132. Thp1 cells were deal with 20 ng/mL “PMA, Phorbol 12-myristate 13-acetate “for 16 hours before LPS stimulation. The ‘rested’ and ‘stimulated’ cells were then prepared for “WCP, whole cell proteome”, “P, phosphoproteome”, “Ac, acetylome” and “Ub, ubiquitinome” analyses. The different color of cell, protein and peptide represent light, medium and heavy SILAC label. Abbreviation: “HPRP, High pH reversed-phase chromatography”; “LC-MS, Liquid chromatography-tandem mass spectrometry”;

### High-confidence integrative proteomics analysis of LPS-stimulated macrophages

We analyzed the overlap of proteins identified by three types of PTM omics and WCP to determine whether the target proteins of different PTMs differed. As shown in **Figure 2A**, approximately half of the proteins identified using WCP did not contain any PTM sites (2554 proteins in Raw cells and 2952 proteins in Thp1 cells). In contrast, 384 proteins in Raw cells and 306 proteins in Thp1 cells carried all three types of PTMs. The table shown in **Figure 2B** provides an overview of the proteins and PTM sites quantified in our study. We quantified 6333 proteins in Raw cells and 6431 proteins in Thp1 cells, 2450 acetylation sites in 1284 proteins in Raw cells and 2183 acetylation sites in 1089 proteins in Thp1 cells, 17,034 phosphorylation sites in 4955 proteins in Raw cells and 18,018 phosphorylation sites in 5162 proteins in Thp1 cells and 7836 ubiquitination sites in 2898 proteins in Raw cells and 7326 ubiquitination sites in 2735 proteins in Thp1 cells (Table S1). The UpSet plots shown in **Figures 2C** and S1C were used to visualize the overlap between PTM sites and proteins in 3 repeated experiments in a matrix layout, allowing us to easily validate the repeatability of the experiments. Pearson’s correlation coefficients for pair-wise comparisons of the log2 M/L abundances were up to 0.96 in Raw cells and 0.88 in Thp1 cells (**Figures 2D** and S1D). Based on these results, our experiments displayed a reliable reproducibility between biological samples.

**Figure 2.**
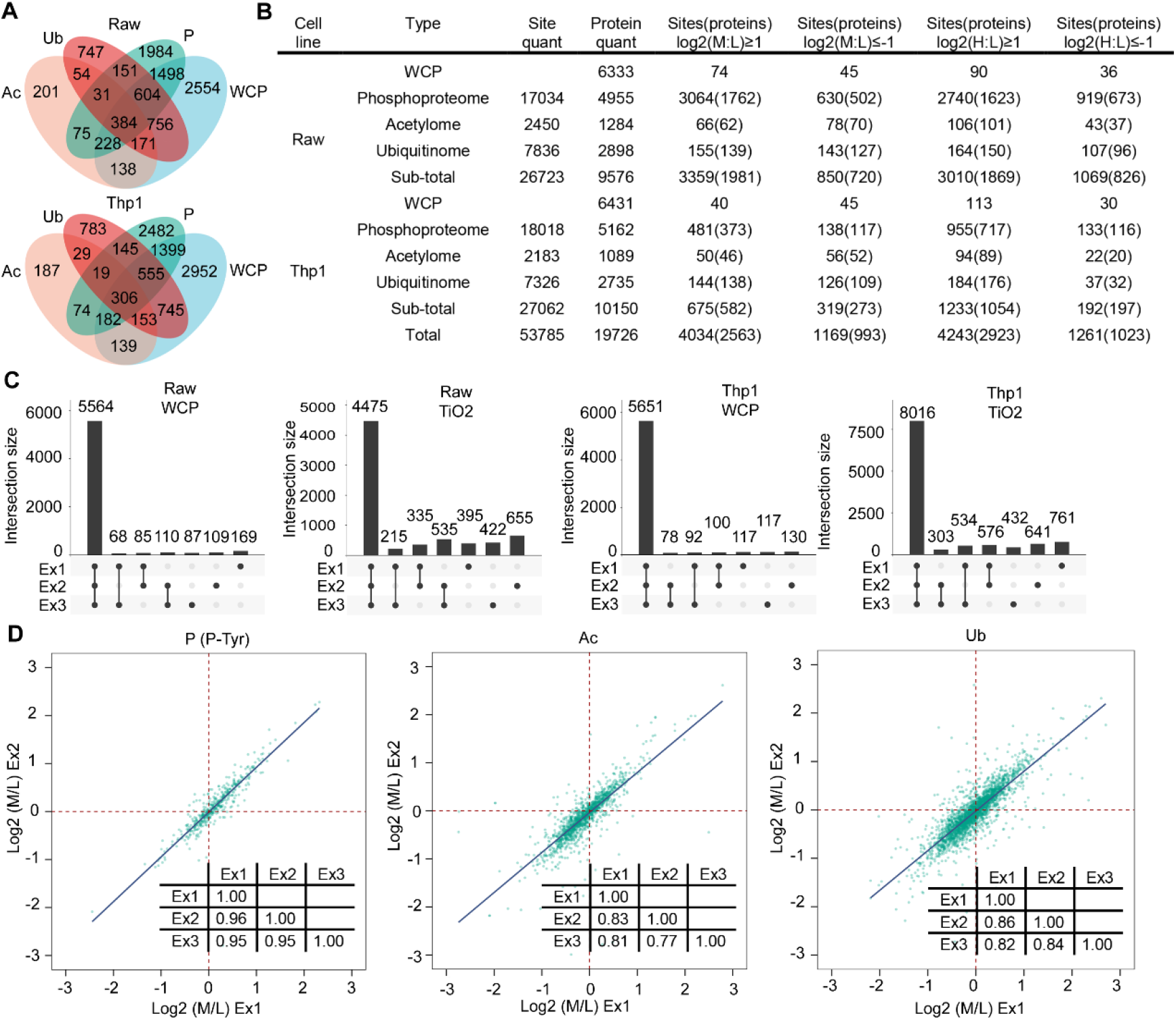
A high-confidence map of integrative proteomics data from LPS-stimulated macrophages. (**A**) Venn diagrams of proteins quantified in the WCP or sites of corresponding proteins quantified in “PTM, posttranslational modification” proteomics. (**B**) Sites and proteins quantified in PTM proteomics and WCP from the two cell types are shown. (**C**) Number of proteins and sites that are quantified in each of 1, 2 or 3 WCP and TiO_2_ experiments in the two cell types. **(D)** Pearson’s correlation plots for two representative experiments (Ex) analysing P-Tyr, Ac and Ub in Raw cells. The inserted table shows Pearson’s correlation coefficients for all three biological duplicates.

### Properties and differences among different types of PTM proteomics

The abundance of different types of PTM on the same protein differed. As shown in **Figure 3A**, phosphorylated proteins usually contained more than one phosphorylation site (61% in Raw cells and 60% in Thp1 cells). The percentage of multiple PTM sites in phosphorylated proteins was higher than in acetylated (39% in Raw cells and 40% in Thp1 cells) and ubiquitinated proteins (55% in Raw cells and 55% in Thp1 cells). We defined the PTM sites with an average 2-fold change (up or down) in three biological duplicates during LPS stimulation as regulated PTM sites. Specifically, approximately 1500 proteins in Raw cells and approximately 700 proteins in Thp1 cells only contained one regulated PTM sites, accounting for larger proportion than other groups (**Figure 3B**). Furthermore, no proteins contained more than two regulated acetylation sites (Figure S2A). Meanwhile, the analysis of the dynamic changes in PTMs is one of interesting part of our research. According to the changes in PTM sites observed in 0.5 hours and 2 hours, we classified the regulated PTM sites into six clusters using the fuzzy c-means method. First, we divided the regulated PTM sites according to the speed at which changes occurred: PTM sites that did not change in 0.5 hours were included in the slow-change group, while the PTM sites that changed were the fast-change group. Then, we divided the fast-change group into lasting and transient groups according to persistence of the change in 2 hours. Finally, we separated the groups into upregulated and downregulated groups, resulting in six clusters. The trends of changes in PTMs are displayed in line charts (**Figures 3C** and S2B). The proportions of “slow” and “fast” items varied in different PTM types (**Figures 3D** and S2C). We defined the proteins with a 2-fold change (up or down) in the PTM sites during LPS stimulation as regulated proteins. Besides, proteins were classified as single-regulated and multi-regulated groups according to the number of regulated PTM sites. In this study, the term “synchrony” was used to refer to regulated proteins in which the regulated PTM sites were in the same slow or fast group mentioned above. In contrast, the term “heterochrony” was used to describe the PTMs of certain proteins that occurred at different speeds. **Figure 3E** provides the summary statistics for proteins classified using our standard. Using the Raw cells as an example, a large proportion (67.4%) of the proteins detected did not contain any regulated PTM sites, while approximately 19.3% of proteins contained a single regulated PTM site. Of the 13.3% proteins containing multiple regulated PTM sites, 41% of them were synchronous, and 59% were heterochronous. Moreover, 13% and 22% of the synchronous proteins and heterochronous proteins contained multi-type of PTM.

**Figure 3.**
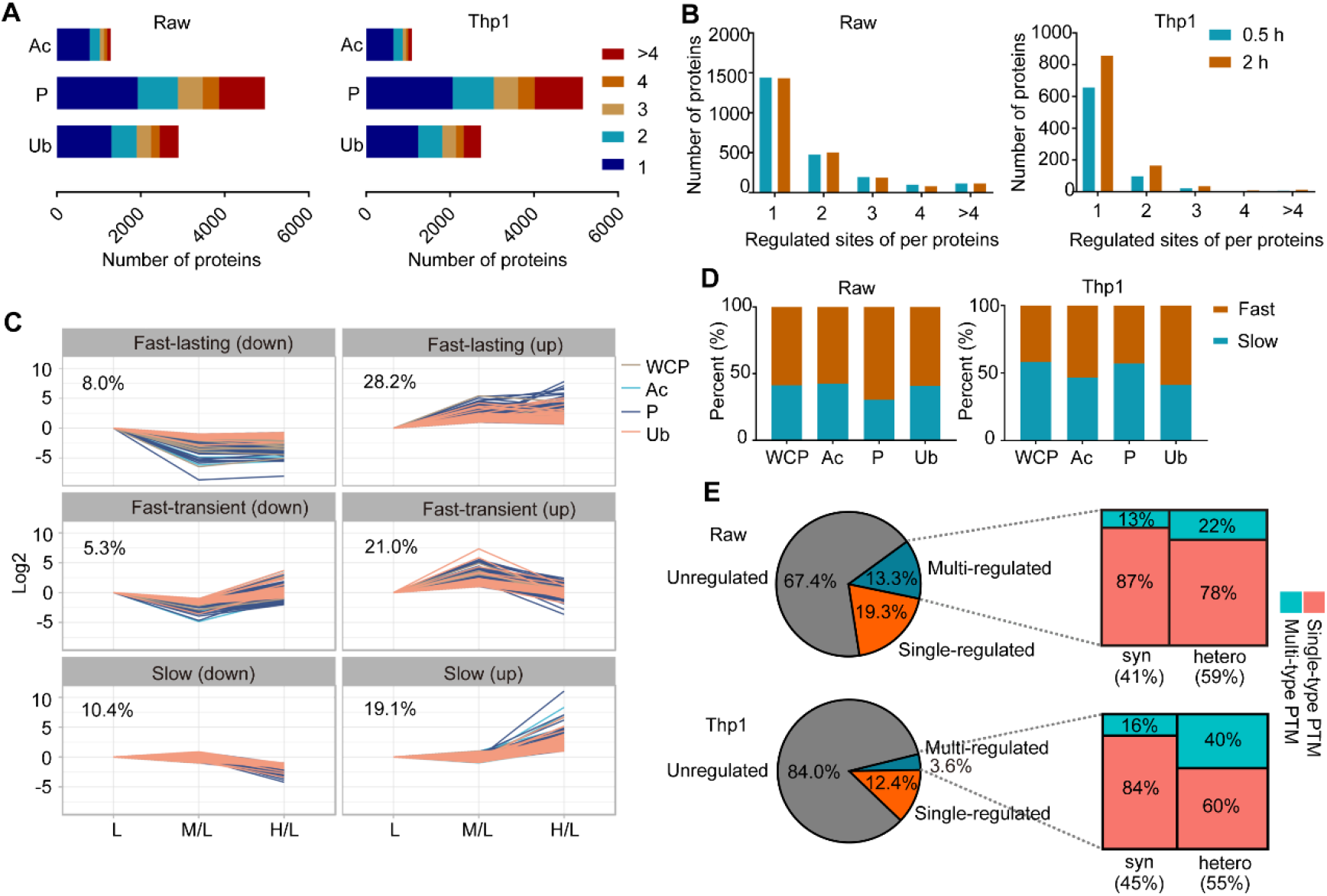
Properties and differences among various types of PTM proteomics. (**A**) Distribution of the number of quantified sites detected in the identical protein in Raw and Thp1 cells. (**B**) Distribution of all regulated PTM sites per protein in macrophages stimulated with LPS for 0.5 and 2 hours. (**C**) Changes in integrative proteomics data in LPS treated Raw cells over time. Regulated proteins and PTM sites were clustered into the six indicated categories using the fuzzy c-means method. (**D**) The distribution of categories of regulated proteins and PTM sites from the integrative proteomics analysis. The six categories in C were combined into two categories according to the speed of change. (**E**) Proteins were classified as unregulated, single-regulated, and multi-regulated groups according to the number of regulated PTM sites. The multi-regulated proteins were classified as “syn, synchronous” and “hetero, heterochronous”, depending on whether all the regulated sites had the same classification in D.

### PTM crosstalk within and across proteins

The density gradient plots shown below illustrate the overall distribution of changes in PTMs. More upregulated PTM sites were identified than downregulated sites in all types of PTMs in both cell lines (**Figures 4A** and S3A). Further, the heatmap of regulated proteins with the three types of PTM and WCP is shown in **Figures 4B** and S3B. We performed an iceLogo analysis to visualize the conserved patterns of protein sequence difference between regulated and unregulated PTM sites (**Figures 4C** and S3C). With the help of the STRING database, we also visualized the comprehensive protein-protein interactions between the regulated proteins (**Figures 4D** and S3D; Table S2). PTM Crosstalk across proteins was based on the assumption that if there is an interaction between two proteins, there is a higher probability that the PTM sites on these two proteins existed crosstalk, especially, when this PTM sites were both regulated in a biological process. Phosphorylation and acetylation mainly targeted proteins related to chromatin assembly and mRNA processing, while ubiquitination mainly targeted proteins related to protein stability (Figure S4). Most of the proteins we identified interacted with other proteins to some extent, indicating the complicated crosstalk among PTMs from different proteins. As shown in the violin plot presented in **Figure 5A**, the average sequence distance of PTM sites in synchronous proteins was closer than in the unregulated proteins. Here, we further divided the sequence distance of PTM sites in heterochronous proteins into two clusters: one only measures the distance between the synchronous PTM sites, while the other only measures the distance between heterochronous PTM sites. The average sequence distances of PTM sites in the two heterochronous groups were much farther than in unregulated proteins and synchronous proteins, and the sequence distance of the synchrony cluster of heterochronous proteins was farther than heterochrony cluster. After counting the numbers of interacting proteins in each group mentioned in Figure 3E, the numbers of interacting proteins in descending order was heterochronous proteins, synchronous proteins, single regulated proteins, and unregulated proteins (**Figure 5B**; Table S3). Additionally, the numbers of total PTM sites in proteins followed the same order as the numbers of interacting proteins (**Figure 5C**). The M/L values of PTM sites in unregulated proteins, single regulated proteins, synchronous proteins and heterochronous proteins was undifferentiated (Figure S5A). We drew a table to present the relationships of different PTMs from the “fast” and “slow” groups. Most of the “fast” type of acetylation sites was more likely to coexist with the “slow” type of phosphorylation sites and acetylation sites, while most of the “fast” type of ubiquitination sites was grouped with the “slow” type of phosphorylation sites and ubiquitination sites. A striking observation from the values presented in this table is that the counts for the intersection of “fast” and “slow” type of phosphorylation sites were particularly large (**Figure 5D**). Those propensities of PTM sites between “fast” and “slow” groups have examples in the immune response. In the *TLR4* pathway, proteins such as *MAP2K4, MAP3K20, NFKB2, TAB3* and *NFKBIE* contained both “fast” phosphorylation sites and “slow” phosphorylation sites. Moreover, *MAPK6* possessed both a “fast” ubiquitination site and a “slow” ubiquitination site, while *TAB2* contained both a “fast” phosphorylation site and a “slow” ubiquitination site (**Figures 5E** and S5B).

**Figure 4.**
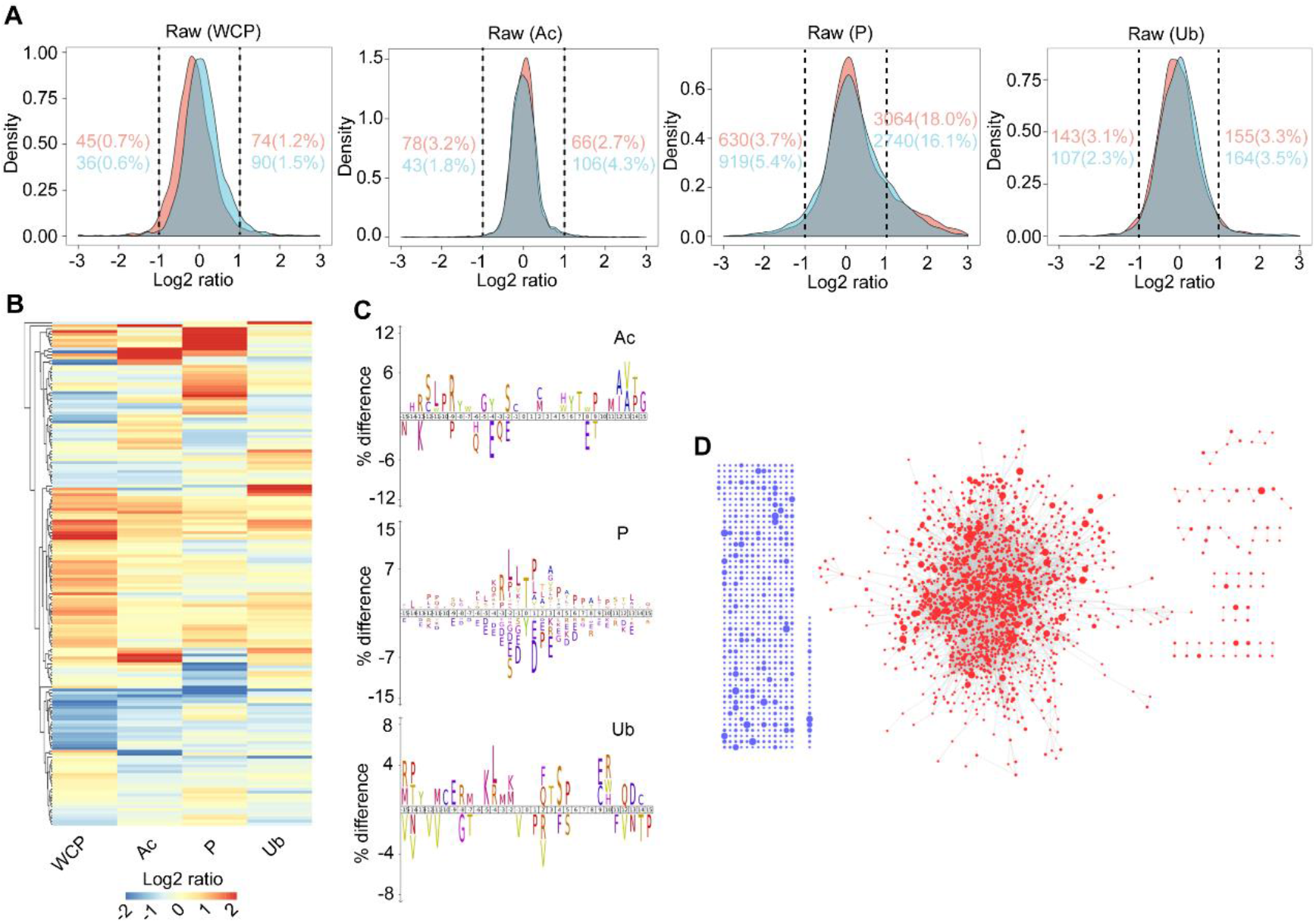
Crosstalk between PTM sites on the different proteins. (**A**) Density gradient diagram of the Log_2_ ratio of proteins and PTM sites in the different proteomes of Raw cells. Carmine and cyan represent 0.5 hours and 2 hours after LPS stimulation, respectively. Carmine and cyan numbers on the left and right represent the number and percentage of regulated proteins and PTM sites in the two time points, respectively. (**B**) Heatmap representation of the Log_2_ (M/L) abundance of proteins quantified in both the WCP and all PTM proteomics in Raw cells. Only proteins with a Log_2_ (M/L) value ≥ 1 or ≤ −1 are shown, and the colour of proteins in PTM proteomics indicate the mean Log_2_ (M/L) ratio of all PTM sites in the protein. (**C**) The iceLogo plots show the difference of amino acid frequency at positions flanking the PTM sites for LPS-regulated PTM sites compared to unregulated PTM sites with a p value ≤ 0.05 in Raw cells. (**D**) Interaction network for proteins with regulated PTM sites in Raw cells. Blue dots indicate the proteins with no interacting partner, while red dots indicate the proteins interacting with other proteins. The size of the dot indicates the number of regulated PTM sites.

**Figure 5.**
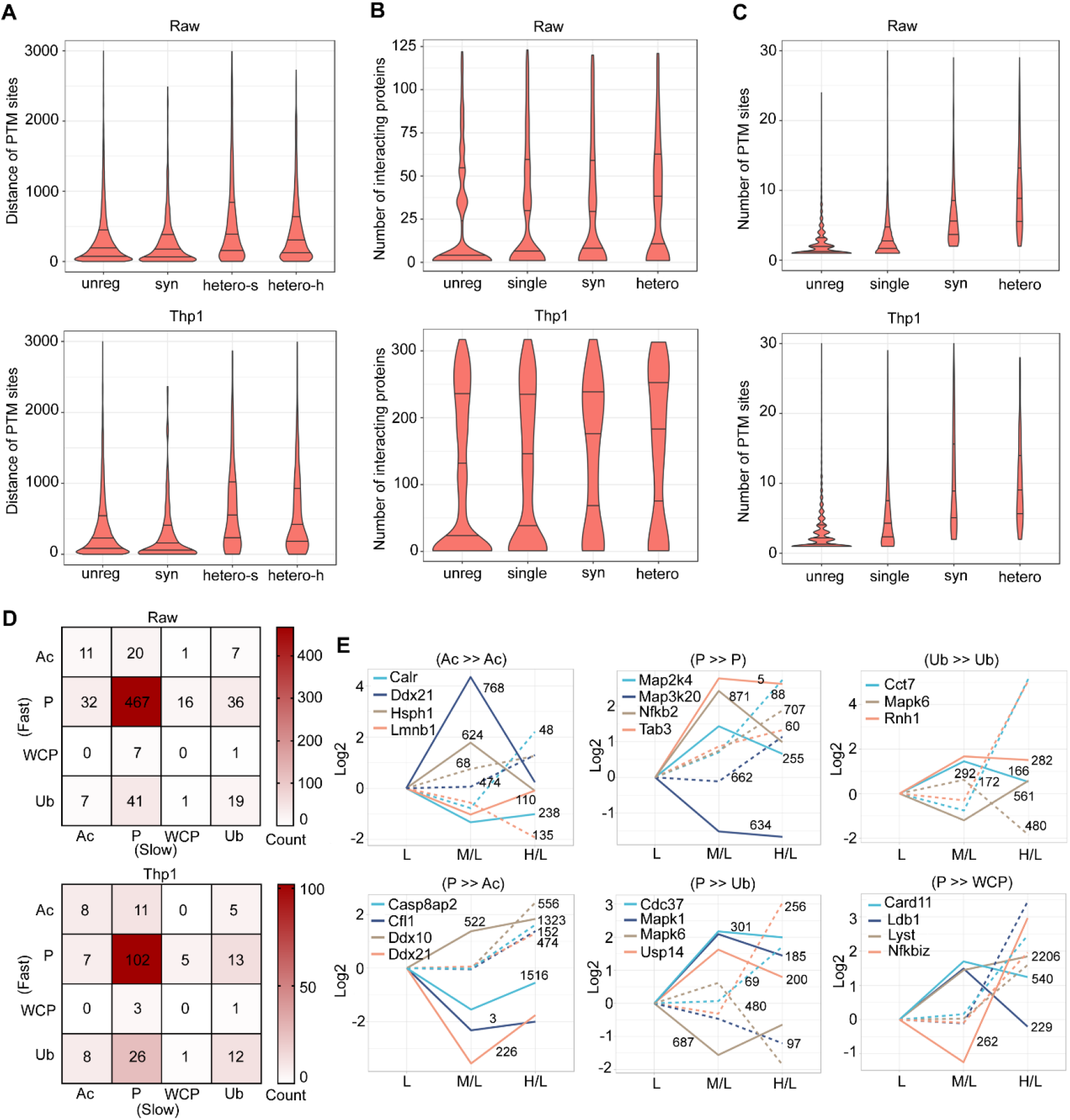
Crosstalk between multiple PTM sites on the same protein. (**A**) Distribution of the sequence distance between different PTM sites for proteins in the “unreg, unregulated”, syn, “hetero-s, the syn sites in hetero proteins” and “heteroh, the hetero sites in hetero proteins” groups. (**B**) Distribution of the number of interacting proteins for proteins in the unreg, “single, single-regulated”, syn and hetero groups. The number of interacting proteins was acquired from IntAct. (**C**) Distribution of the number of identified PTM sites for proteins in the unreg, single, syn and hetero groups. (**D**) Heatmap indicating all relationships between different PTM types and the WCP. (**E**) Selected proteins in Raw cells belonging to the important combination types listed in D. The solid line represents a ‘fast’ regulated event and the dotted line represents a ‘slow’ regulated event. The number next to the line represents the site of PTM on the corresponding protein. The PTM in the left of “≫” is ‘slow’ PTM and The PTM in the right of “≪” is ‘fast’ PTM. For A, B and C, the lower, median and upper lines in each violin plot correspond to25%, 50% and 75%, respectively.

### LPS induces both degradative and non-degradative ubiquitination

Ubiquitylation usually results in two distinct outcomes: degradative and non-degradative processes. We integrated WCP and ubiquitinome datasets acquired from cells treated with or without MG132 to clearly distinguish the results of ubiquitination after LPS stimulation. We first investigated whether the number of proteins would be influenced by the level of mRNA. The levels of several inflammatory cytokines, namely, *IL-1, IL-16, IL-18, TGFB1* and *TNF*, which should increase after LPS stimulation, did not change in both Raw and Thp1 cells after 0.5 hours of stimulation (**Figure 6A**). Meanwhile, real-time qPCR revealed that the mRNA levels of the proteins whose levels were significantly regulated in our WCP datasets did not increase (**Figure 6B**). Together, the level of mRNA had little effect on the quantity of proteins at 0.5 hours after LPS stimulation. The relative abundance of ubiquitin lysine sites indicated that the ubiquitination of K48 (usually results in degradation) dominated, followed by the ubiquitination of K63 (usually does not result in degradation) (**Figure 6C**). Additionally, the ubiquitination of K48 and K63 was relatively stable during LPS stimulation (Figure S6A). A closer inspection of the ubiquitinome and WCP revealed that some ubiquitination sites (4% in Raw cells and 3% in Thp1 cells) and proteins (1% in both Raw and Thp1 cells) in LPS-stimulated macrophages dramatically affected by MG132 (**Figures 6D** and S6B). We next predicted the outcomes of ubiquitination proteins according to the hypothesis that changes in ubiquitination are inversely proportional to changes in the levels of proteins for degradative ubiquitination (**Figure 6E**). Notably, non-degradative ubiquitination was widespread after LPS stimulation, including *NFKB1, MAPK1, TRAF1* and other proteins (**Figure 6F**). Notably, MG132 modulated LPS-induced inflammation (Figure S6C).

**Figure 6.**
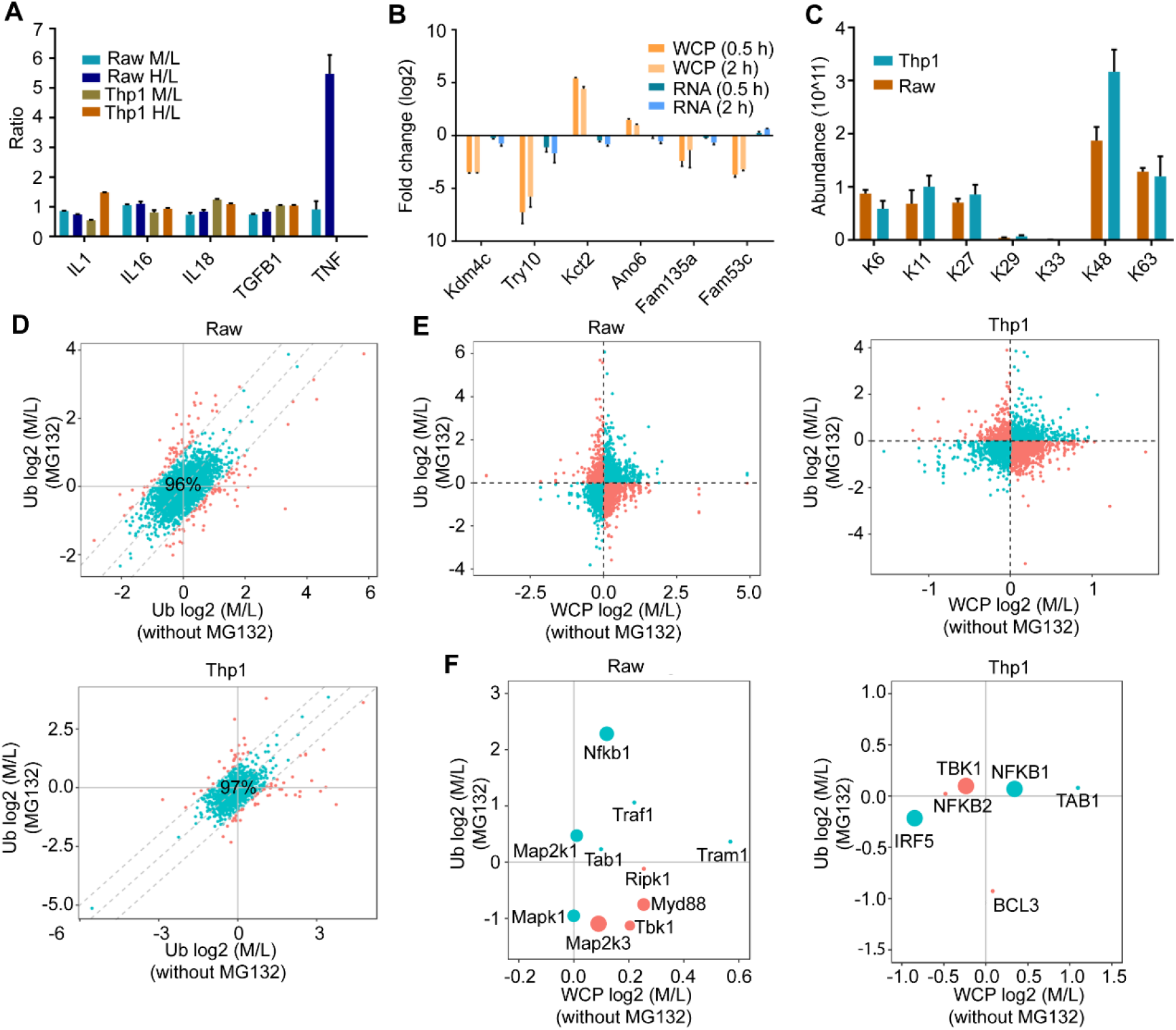
Degradative and non-degradative ubiquitination both exist in the LPS-stimulated ubiquitinome. (**A**) WCP protein ratios for selected inflammatory factors from three biological replicates are shown. (**B**) The levels of proteins and RNAs (Fold changes related to 0 h) for selected genes in Raw cells with LPS stimulation from three biological replicates are shown. (**C**) Abundance of ubiquitin lysine sites quantified in the Ub of LPS-stimulated macrophages. (**D**) Comparison of Log_2_ (M/L) value of the ubiquitination site abundance from the LPS-stimulated Ub of cells treated with or without MG132 for 2 hours. Sites that exhibited a ≥ 1 Log_2_ (M/L) difference in untreated and MG132-treated cells were considered dramatically affected by MG132 (carmine). (**E**) Comparison of Log_2_ (M/L) values in the WCP of cells treated without MG132 and the Ub of cells treated with MG132. Sites showing the same changes in the two types of omics data listed above are predicted to be non-degradative ubiquitinated sites (cyan), while the remaining sites are degradative ubiquitinated sites (carmine). (**F**) Proteins in the TLR4 pathway are used to illustrate the classification in E. For A, B and C, error bars represent the standard error of the mean.

### New proteins and PTMs involved in inflammatory responses

We integrated the WCP, acetylome, phosphoproteome, and ubiquitinome datasets to construct a network of PTM patterns for regulated proteins in the *MAPK* and *NFKB* signalling pathways and to illuminate the PTMs of proteins in major inflammatory signalling pathways that are activated by LPS stimulation. We mapped protein, acetylation, phosphorylation and ubiquitination level ratios at different times in this network. Twenty-two proteins *(IRAK2, IRF3, TAB2, TAB3, TBK1, TRIF, MAPK9, MAPK11, MAPK12, MAP2K3, MAP2K4, MAP2K7, MAPKAPK2, MNK2, MSK1, MSK2, C-Rel, NFKB1, NFKB2, NFKBIE, NFKBID*, and *NFKBIZ*) in this network exhibited upregulated phosphorylation sites, while four proteins (*TAB3*, *MAP2K7*, *MSK2*, and *NFKBIZ*) in this network displayed downregulated phosphorylation sites. Besides, some of these proteins contained more than one regulated phosphorylation sites, for example, *TAB3* contained seven regulated phosphorylation sites but only phosphorylation in S60 been well studied. Interestingly, *TAB3* and *MAP2K7* contained both upregulated and downregulated phosphorylation sites. Acetylation and ubiquitination of proteins in this network were generally unregulated, except the acetylation of K310 in *RelA*. Nevertheless, two proteins (*MYD88* and *MAPKAPK2*) contained low levels of regulated (log2 ratio ≥ 0.5 or ≤ −0.5) ubiquitination sites (**Figure 7A**). Regulated PTM sites widely existed in several protein families *(CASPASE* family, *MAPK* family, *MAP2K* family, *MAP3K* family, interferon related family, *NLRP* family, *TRAF* family, and *RIPK* family) associated with inflammation in both Raw and Thp1 cells (**Figures 7B** and S7). All the PTM data obtained in this study were visualized in a PTM-inflammation website (**Figure 8A**). To help identify new proteins that involved in the inflammatory response, we filtered 95 proteins that contained significantly regulated PTM sites but lacked studies on inflammation. We further screened these proteins using corresponding siRNAs, and the results are displayed in **Figure 8B** and Table S4. Homology comparison of regulated proteins identified in Raw264.7 and Thp1 cells revealed that the post-translational modifications of 91 proteins were conserved between humans and mice (Table S5).

**Figure 7.**
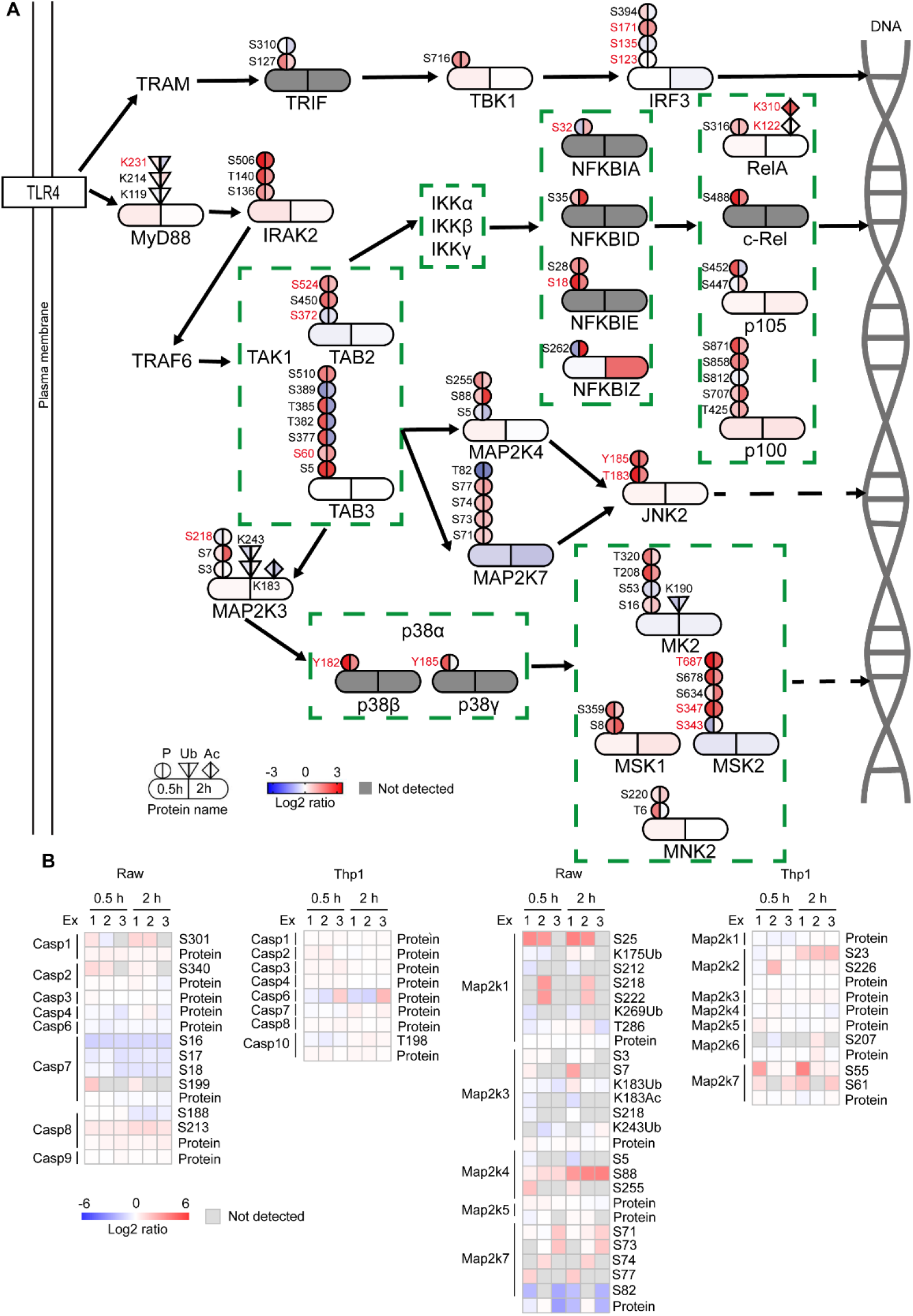
New PTMs in TLR4 signaling pathway involved in inflammatory responses. (**A**) Regulated proteins within the *TLR4* pathway are grouped by function, and arrows indicate the direction of signal transduction. The colours on the left and right represent Log_2_ ratios of the indicated proteomics datasets from cells stimulated with LPS for 0.5 hours and 2 hours, respectively. PTM sites with known functions based on UniProt and previous articles are coloured in red [31–41]. (**B**) Heatmap representation of the intensity of proteins and PTM sites involved in inflammatory signalling pathways detected in cells stimulated with LPS for 0.5 hours and 2 hours.

**Figure 8.**
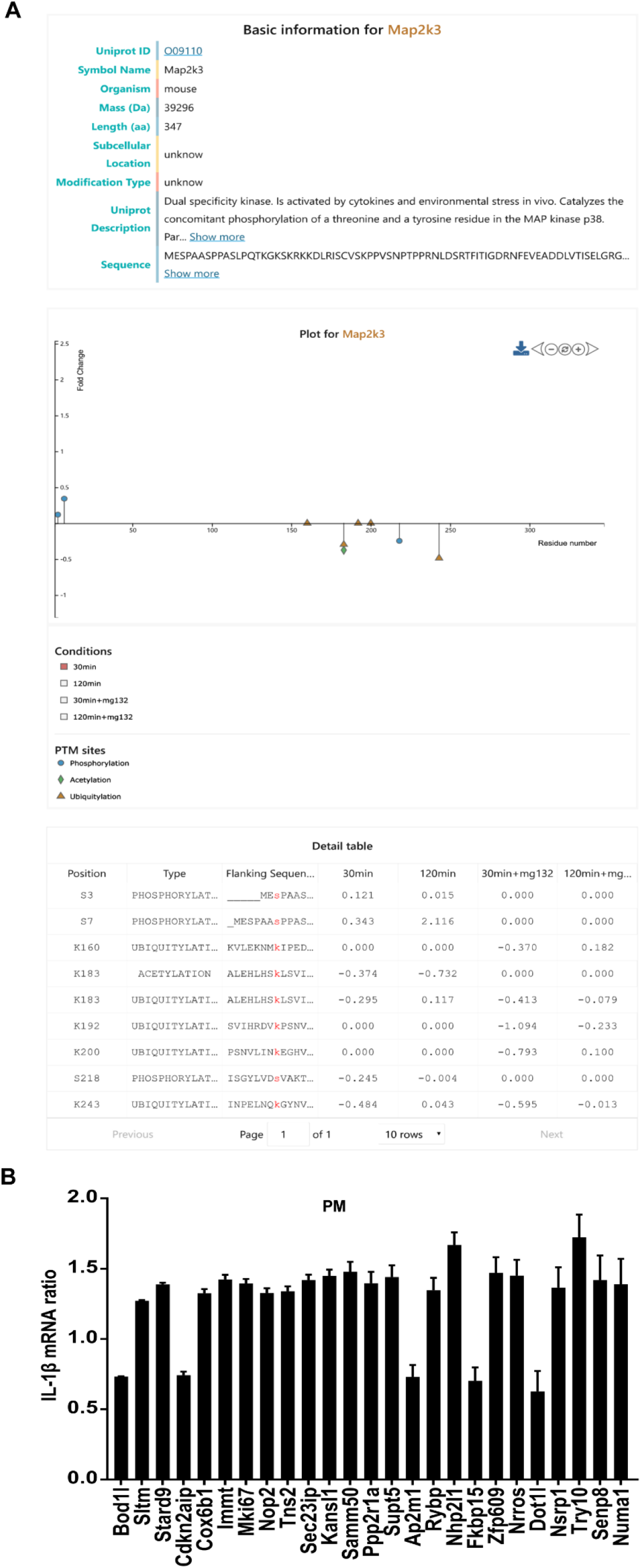
New proteins involved in inflammatory responses. (**A**) *Map2k3* served as an example of the PTM-inflammation website (http://ptm-inflammation.cn). The figure shows basic information about the protein, such as the sub-cellular location, modification type, UniProt description, sequence, etc. It also includes a visualization of post-translational data and a table providing detailed data. (**B**) The results of the siRNA screen for proteins containing apparently regulated PTM sites in mouse “PM, peritoneal macrophage” cells. The ratio of the *IL-1β* mRNA was generated by comparing the effect of a certain siRNA with a negative control siRNA. Figures were based on data obtained from two biological replicates each containing three technical replicates, and error bars represent the standard error of the mean.

## Discussion

Previous studies have reported LPS-induced proteomics or phosphoproteomics data in Raw264.7 and Thp1 cells [19, 20]. However, these studies focused on functional analysis of differential proteins and the type of post-translational modification was primarily phosphorylation. In this study, we integrated WCP, acetylome, phosphoproteome and ubiquitinome datasets to identify LPS stimulation-dependent PTM events in macrophages. These data also provided insights into the crosstalk between PTMs during the inflammatory response. Although we didn’t validate the PTM data by biochemical approaches, our results were consistent with previous findings, such as the phosphorylation on Y182 of *P38β*, the phosphorylation on Y185 of *P38γ* and the phosphorylation on S18 of *NFKBIE* (Figure 7A). Besides, we performed siRNA experiments in mouse PM. As expected, the inflammatory response was affected by reducing the proteins whose PTMs were significantly altered in cell lines.

Among those datasets, phosphorylation had a significant advantage in terms of the number of quantified sites. For all PTMs in both types of macrophages, except phosphorylation in Raw cells, the proportion of upregulated sites exceeded the proportion of downregulated sites, and the proportion of upregulated sites increased while the proportion of downregulated sites decreased as the period of LPS stimulation increased. We believe that the faster reaction rate of phosphorylation leads to the different trend of phosphorylation in Raw cells. Additionally, the PTM properties of different cells were slightly different, and different PTMs usually occurred in the proteins of distinct functional groups. However, the WCP showed obvious randomness in functional groups, suggesting that the LPS-induced signal transduction was less dependent on the changes in protein levels.

The time series analysis has increasingly been applied to proteomics [21, 22]. In our study, a large number of proteins contained at least one type of regulated PTM. Within one protein, crosstalk events usually occurr among nearby PTM sites, so the sequence distances between crosstalk pairs are shorter than average [23]. Therefore, the regulated PTM sites in proteins that only contain synchronously regulated PTM sites are more likely to be involved in crosstalk due to their shorter sequence distance than unregulated PTM sites. However, the sequence distance for both synchronously and heterochronously regulated PTM sites in proteins that contain heterochronously regulated PTM sites was farther than unregulated PTM sites. Additionally, more interacting partners was observed for proteins containing heterochronously regulated PTM sites than proteins only containing synchronously regulated PTM sites. Overall, we concluded that the regulated PTM sites in proteins containing heterochronously regulated PTM sites are more likely to have crosstalk with PTM sites in the corresponding interacting proteins. It can thus be suggested that the PTM sites participating in crosstalk between proteins have farther sequence distance because they tend to located in different domains in a protein. Interestingly, the number of PTM sites was proportional to the number of interacting proteins.

MG132, which inhibits the degradation of ubiquitinated protein, has been widely used to study the molecular mechanism of inflammation [24–26]. However, MG132 could directly influence the ubiquitination of certain proteins or indirectly influence ubiquitination by inhibiting the inflammatory response of macrophages. Thus, it is difficult to distinguish degradative and non-degradative ubiquitination by comparing the outcomes in cells treated with or without MG132. Therefore, we proposed a more reliable method to determine the levels of degradative and non-degradative ubiquitination according to whether the changes in protein levels were consistent with the changes in ubiquitination levels after the addition of MG132. Using this technique, we discovered both degradative and non-degradative ubiquitination are prevalent in the immune response to LPS stimulation. The premise of this method is to limit the reaction time, so that the effect of gene expression on protein levels can be ignored. Complementarily, the integrated analysis of ubiquitinome, WCP and transcriptome data solves this problem when the effect of gene expression on protein level is unable to be ignored [27]. Although these methods eliminate the negative effects of MG132, a specific inhibitor of deubiquitination for a specific protein is preferred.

In conclusion, this study provides high-quality datasets of multiple PTMs to serve as a resource for screening new molecules involved in the inflammatory response. The identified PTM sites in the known inflammation-related proteins could help to discover the underlying molecular mechanisms in inflammatory response. Importantly, our in-depth analyses have identified the interrelationships among the most common PTMs and pave the way for future studies aiming to determine the relationships among other PTMs.

## Materials and methods

### Reagents

MG-132 (Selleck, Cat# S2619, Houston, USA), PR-619 (Selleck, Cat# S7130, Houston, USA), trypsin (Beierli, Cat# BELT001), PTMScan ubiquitin remnant motif kit (Cell Signaling Technology, Cat# 5562, Bossdun, USA), PTMScan acetyl-lysine motif kit (Cell Signaling Technology, Cat# 13416, Bossdun, USA), PTMScan phospho-tyrosine rabbit mAb kit (Cell Signaling Technology, Cat# 8803, Bossdun, USA), high-select Fe-NTA phosphopeptide enrichment kit (ThermoFisher, Cat# A32992, Waltham, USA), high-select TiO_2_ phosphopeptide enrichment kit (ThermoFisher, Cat# A32993, Waltham, USA), protease and phosphatase inhibitor cocktail (Beyotime, Cat# P1049), DMEM for SILAC (Gibco, Cat# 88364), dialyzed fetal bovine serum (Gibco, Cat# 30067334), L-lysine-^13^C_6_-^15^N_2_ (Gibco, Cat# 88209), L-arginine-^13^C_6_-^15^N_4_ (Gibco, Cat# 89990), L-lysine-4,4,5,5-D4 (Gibco, Cat# 88438), L-arginine-^13^C_6_ (Gibco, Cat# 88433), and RPMI 1640 medium for SILAC (Gibco, Cat# 88365).

### Cell culture and sample preparation

Raw264.7 and Thp1 cells were cultured in DMEM or RPMI 1640 medium without lysine and arginine but excess L-proline. Cells grown in light media (supplemented with 100 mg/L L-lysine and 100 mg/L L-arginine), medium media (supplemented with 100 mg/L L-lysine-4,4,5,5-D4 and 100 mg/L L-arginine-^13^C_6_) and heavy media (supplemented with 100 mg/L L-lysine-^13^C_6_-^15^N_2_ and 100 mg/L L-arginine-^13^C_6_-^15^N_4_) were activated with 100 ng/mL LPS for 0, 0.5 and 2 hours, respectively. Before the follow-up experiments, we detected the SILAC experiments with a labeling efficiency of over 98%. For the MG132 experiments, all SILAC-labelled cells were treated with 5 μM MG132 for 2 hours. A 10-cm dish of Cells were harvested for each experiment and lysed with freshly prepared lysis buffer (8 M urea, 150 mM NaCl, 50 mM Tris-HCl (pH 8.0), 1X phosphorylase inhibitor, 1X EDTA, 50 μM PR-619, and 1x protease inhibitor). Protein concentrations were estimated using the BCA kit and lysates from each cell line treated with light, medium and heavy media were combined in a 1:1:1 ratio. The combined lysates were reduced with 10 mM dithiothreitol (DTT), carbamidomethylated with 30 mM iodoacetamide (IAA) and digested overnight with trypsin. Peptides were acidified with trifluoroacetic acid (TFA), desalted with preconditioned C18 Sep-Pak SPE cartridges and lyophilized for 48 hours with a vacuum lyophilizer (LABCONCO, Kansas, USA). Peritoneal macrophages were acquired from 6-8-week-old male C57/BL6 mice. This study was approved by the ethics committees of the First Affiliated Hospital of Zhejiang University.

### Enrichment of ubiquitinated, acetylated and phosphorylated peptides

For proteomics analyses of acetylation, p-Tyr and ubiquitination, lyophilized peptides were resuspended in cold 1× IAP buffer and incubated with corresponding cross-linked antibody beads for 2 hours at 4°C with end-over-end rotation. Beads were softly washed three times with ice-cold 1× IAP buffer and twice with ice-cold MS water. Modified peptides were eluted twice by adding 50 μl of 0.15% TFA. For TiO_2_ and Fe-NTA proteomics, lyophilized peptides were treated using the methods described in the protocols provided with the corresponding kits. The enriched peptides were quickly dried by vacuum centrifugation and divided into eight fractions using high pH reversed-phase chromatography (HPRP). All peptides were desalted using homemade stage tip chromatography and dried by vacuum centrifugation prior to the MS analysis.

### LC-MS/MS Analysis

All peptide samples were resuspended in 2% acetonitrile (ACN) and 0.1% formic acid (FA). Then, peptides were separated by nanoLC-MS/MS using an UltiMate 3000 RSLCnano system (ThermoFisher, Waltham, USA) and analysed using Q Exactive HF-X (ThermoFisher, Waltham, USA). Gradient elution was performed at 32°C over 120 minutes using a gradient of 3–80% ACN in 0.1% FA. The MS spectra of WCP, ubiquitinome, TiO_2_ and Fe-NTA samples were acquired at a resolution of 120,000 with a mass range of 300–1500 m/z and an AGC target of 3E6, while acetylome and p-Tyr spectra were acquired at a resolution of 60,000. MS2 spectra were acquired at a resolution of 30,000, and HCD fragmentation was performed with a collision energy of approximately 27% NCE. The isolation window was set to 1.0 m/z and the dynamic exclusion window was set to 30 s.

### Identification and quantification of the proteome

MaxQuant (version 1.6.2.10) was used for protein identification and quantification. The human and mouse UniProtKB databases were utilized as the search databases. The variable modifications of the acetylome, phosphoproteome and ubiquitinome included oxidation (M), acetyl (protein N-term), and corresponding PTMs. Carbamidomethyl (C) was set as the fixed modification. The maximum number of modifications for a peptide was set to 5. Trypsin was set as the digestion enzyme, and the maximum value of missed cleavage sites was 2. Additionally, we used 20 ppm as the ion tolerance in the first search and 4.5 ppm as the ion tolerance in the main search. Both peptide and protein identification were performed at an FDR < 1%. The default parameters of MaxQuant were adopted if not described above.

### Real-time qPCR and siRNA screening experiment

Raw cells were treated with 100 ng/mL LPS, and RNA was isolated using the RNA extraction kit (Feijie) according to the manufacturer’s protocol. The cDNA templates were synthesized using the RNA reverse transcription kit (Takara, Tokyo, Japan). Quantitative PCR was performed using TB Green^®^ Premix Ex Taq™ II (Takara, Tokyo, Japan). The siRNA was purchased from Dharmacon and transfected into Raw and PM cells using the DharmaFECT 1 siRNA Transfection Reagent.

### Functional Annotation Enrichment Analysis

Proteins containing PTM sites with a fold change ≥2 were selected for functional annotation enrichment analysis using WebGestalt 2019 to identify the pathways in which different PTMs were enriched in cells stimulated with LPS based on the proteomics data [28]. We chose the top three categories of the forty categories visualized in the report for each cell and time point. The heat map was constructed using the pheatmap package in R software.

### The PTM consensus motifs

We analysed sequences within −15 to +15 amino acids of the quantified PTM sites (the PTM site was located at position 0) to explore the consensus sequence of amino acids surrounding each PTM. IceLogo was used to generate the amino acid sequence diagram [29].

### Statistical analysis

The R framework (version 3.5.1) and GraphPad Prism (version 7) software were used to perform all statistical analyses of the bioinformatics data. All the proteomics datasets were performed in biological triplicates. The real-time qPCR analysis of mRNA abundance was performed in biological triplicates. The siRNA screening experiment was performed using biological duplicates. The table listing PTM sites was filtered to remove entries with a localization probability less than 90%. The PTM sites with a membership greater than 0.6 were classified into the corresponding category using the fuzzy c-means method. The interaction network only retained the interactions with a STRING database score greater than 0.8.

### Data availability

The mass spectrometry proteomics data have been deposited to the ProteomeXchange Consortium (http://proteomecentral.proteomexchange.org) via the iProX partner repository [30] with the dataset identifier PXD015527. PTM-inflammation website (http://ptm-inflammation.cn) is freely accessible. All other data supporting our findings are available in the article and supplemental information.

## Authors’ contributions

FY.J, HH.Z, MH.Z and ZY.J performed the majority of experiments. FY.J, YM.L and MH.Z performed the majority of data and statistical analysis. FY.J, HH.Z and MH.Z conceived and designed experiments. FY.J, MH.Z, YM.L and QR.L wrote and edited the manuscript. LJ.L directed the study. XX.OY, SN.Z and T.W assisted with experiments.

## Acknowledgements

We thank zeyu sun and Jing Jiang for mass spectrometry technical support. We thank Danhua Zhu, Xiaopeng Yu and Yanhong Zhan for helpful discussions. This work was supported by the National Key Research and Development Program of China (2016YFC1101304/3), the National Key Research and Development Program (2017YFC1200100), the National Natural Science Foundation of China (81400589), Chinese national science and technology major project of the 13th Five-year plan(2017ZX10202202-001-008), and Science Fund for Creative Research Groups of the National Natural Science Foundation of China (81721091).

## Supplementary material

**Figure S1.**
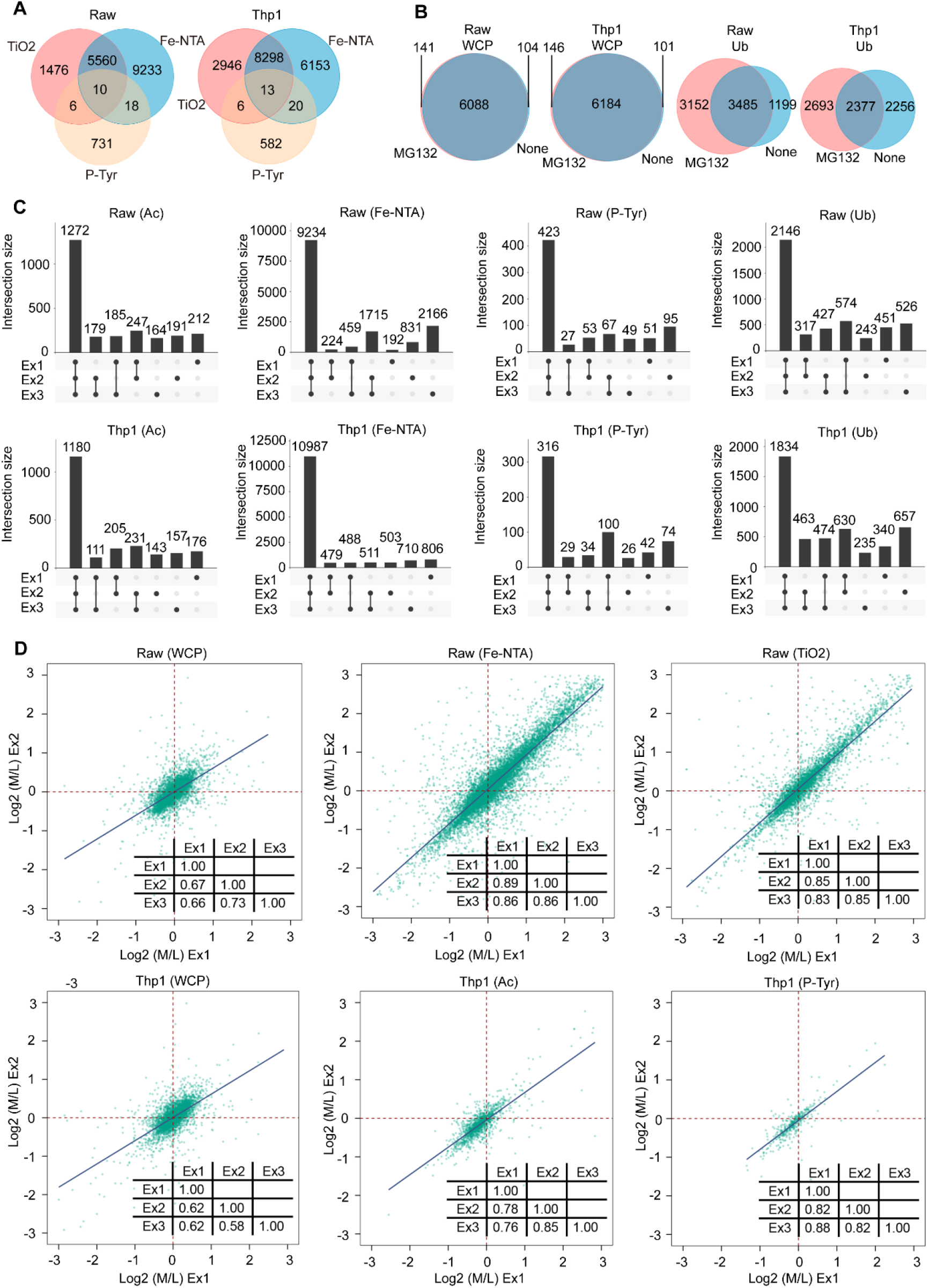
Reproducible integrative proteomics analysis of LPS-stimulated macrophages. (**A**) Intersection of phosphosites quantified with the three methods used to analyse the P. (**B**) Overlap between proteins quantified in the WCP and diGly sites quantified in the Ub in “MG132, presence of MG132” and “None, absence of MG132” groups. (**C**) Number of PTM sites that were quantified in each of 1, 2 or 3 Ac, Fe-NTA, P-Tyr and Ub experiments in two cell types. (**D**) Pearson’s correlation plots for two “Ex, representative experiments” from WCP, Fe-NTA, TiO_2_, Ac and P-Tyr in Raw and Thp1 cells. The inserted table shows Pearson’s correlation coefficients for all three biological duplicates.

**Figure S2.**
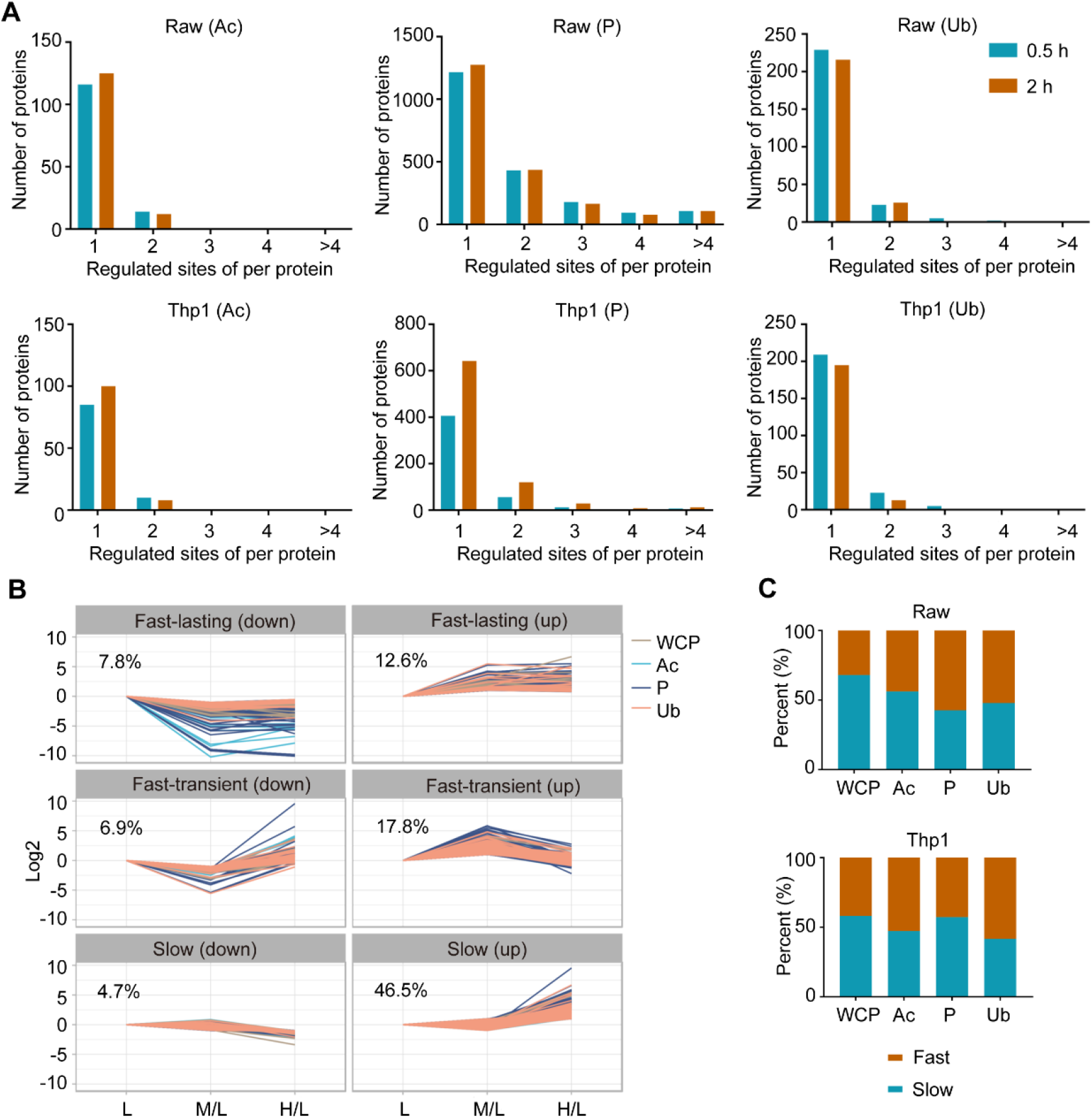
Properties and differences among various types of proteomics approaches. (**A**) Distribution of regulated acetylated, phosphorylated and ubiquitinated sites per protein in macrophages stimulated with LPS for 0.5 and 2 hours. (**B**) Changes in integrative proteomics data obtained from LPS-treated Thp1 cells over time. Regulated proteins and PTM sites were clustered into the six indicated categories using the fuzzy c-means method. **(C)** The distribution of categories of regulated sites in the corresponding proteins identified using integrative proteomics. The six categories in B were combined into two categories according to the speed of change.

**Figure S3.**
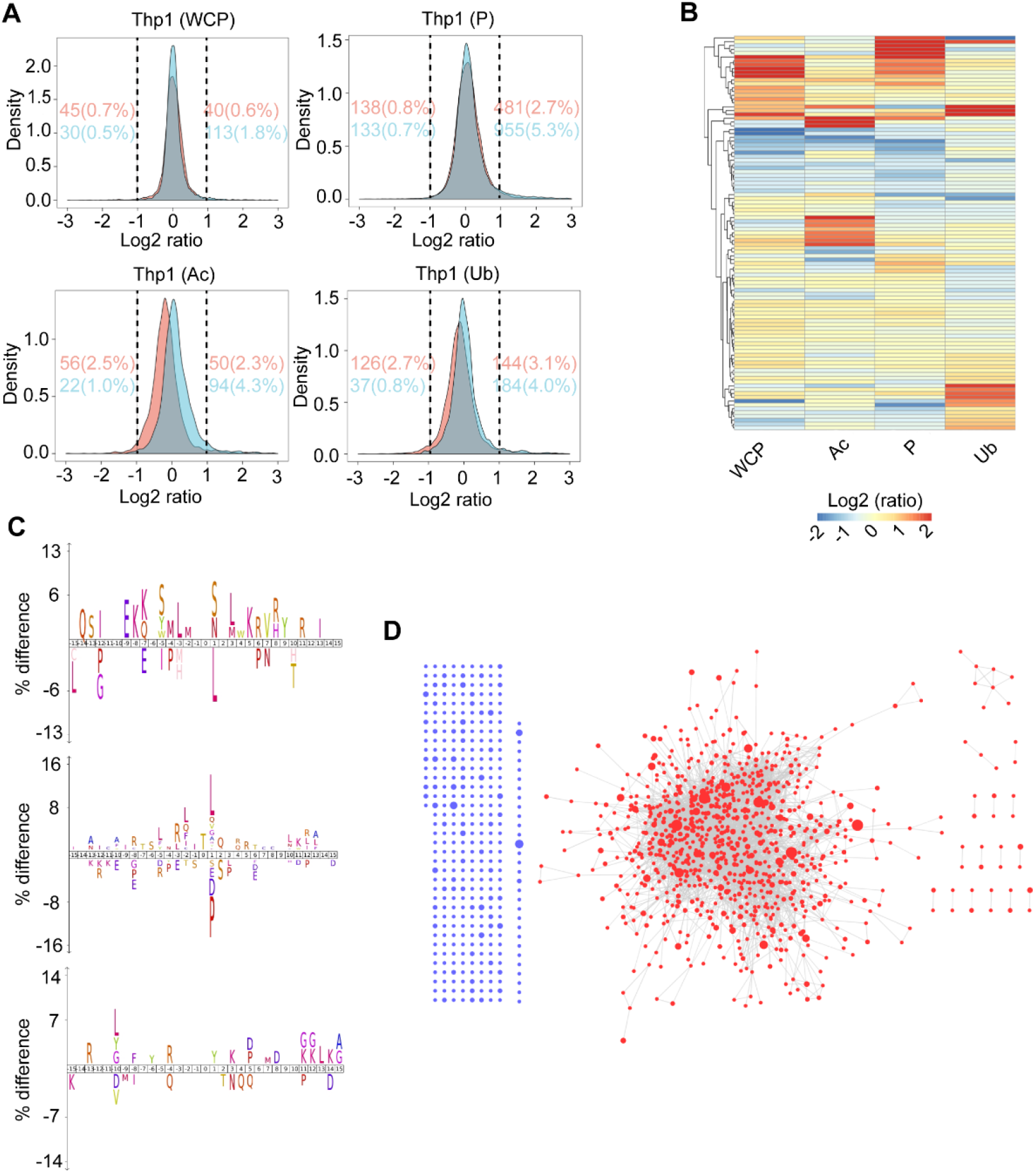
PTM crosstalk between different proteins. (**A**) Density gradient diagram of the Log_2_ ratio of proteins and PTM sites in the different proteomes of Thp1 cells. Carmine and cyan represent cells stimulated with LPS for 0.5 hours and 2 hours, respectively. Carmine and cyan numbers on the left and right represent the number and percentage of regulated proteins and PTM sites in the two time points, respectively. (**B**) Heatmap representation of the Log_2_ (M/L) of the abundance of proteins quantified in Thp1 cells using the WCP and all PTM omics methods. Only proteins with a Log_2_ (M/L) value ≥ 1 or ≤ −1 are shown, and the colour of proteins identified using PTM omics indicate the mean Log_2_ (M/L) ratio of all PTM sites in the protein. (**C**) The iceLogo plots show the difference of amino acid frequency at positions flanking the PTM sites for LPS-regulated PTM sites compared to unregulated PTM sites with a p value ≤ 0.05 in Thp1 cells. (**D**) Interaction network for proteins with regulated PTM sites in Thp1 cells. Blue dots indicate the proteins with no interacting partners, while red dots indicate the interacting proteins. The size of the dot indicates the number of regulated PTM sites.

**Figure S4.**
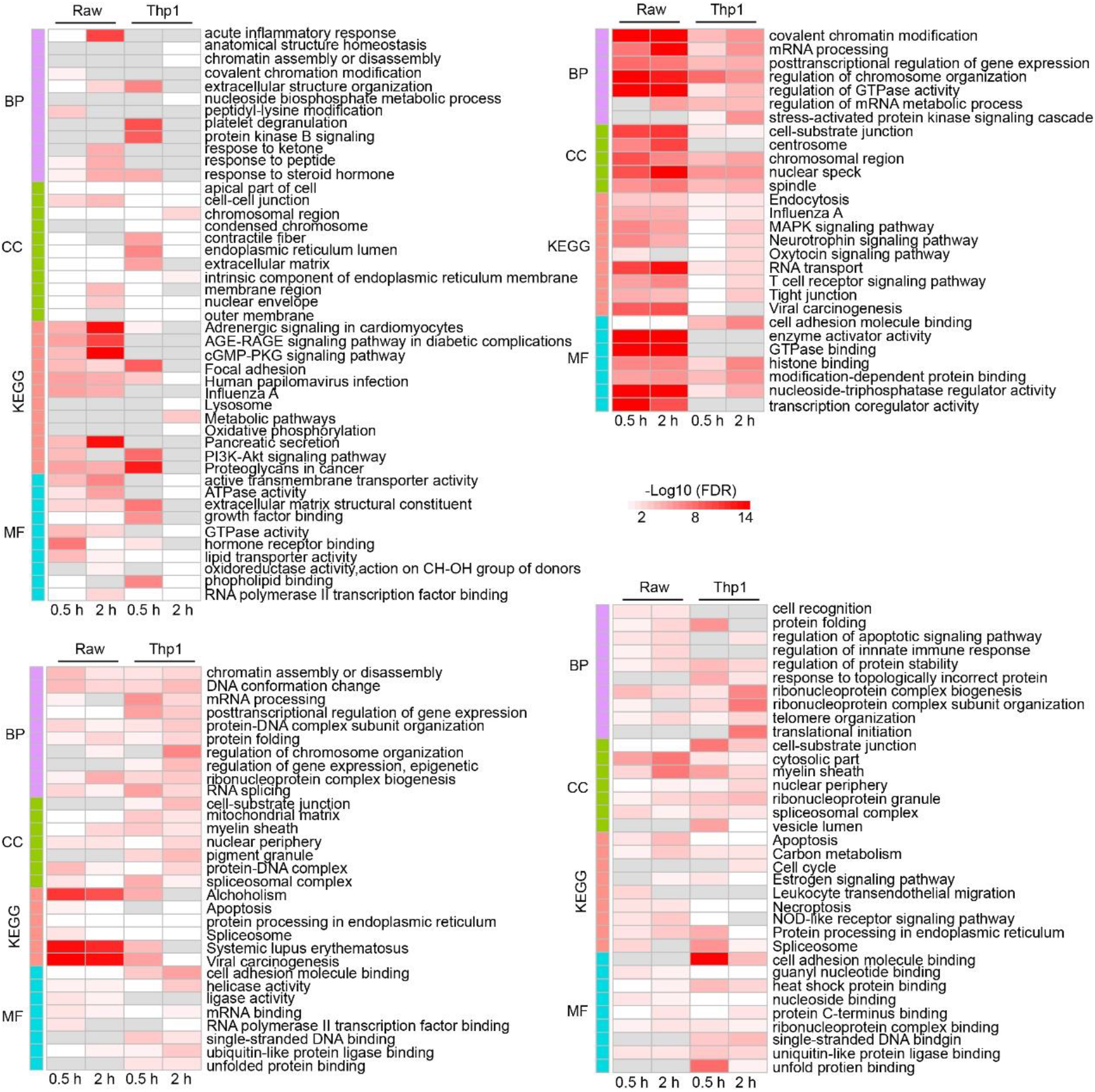
Annotation enrichment analysis of regulated proteins. Annotation enrichment analysis of proteins with regulated expression level (top left panel), proteins with regulated acetylation sites (bottom left panel), proteins with regulated phosphorylated sites (top right panel) and proteins with regulated ubiquitinated sites (bottom right panel). Only the terms with the top three −Log_10_ (FDR) values in each of the following categories for all time points and both cell lines are shown: “BP, biological processes”, “CC, cellular compartments”, “KEGG, pathways” and “MF, molecular functions”.

**Figure S5.**
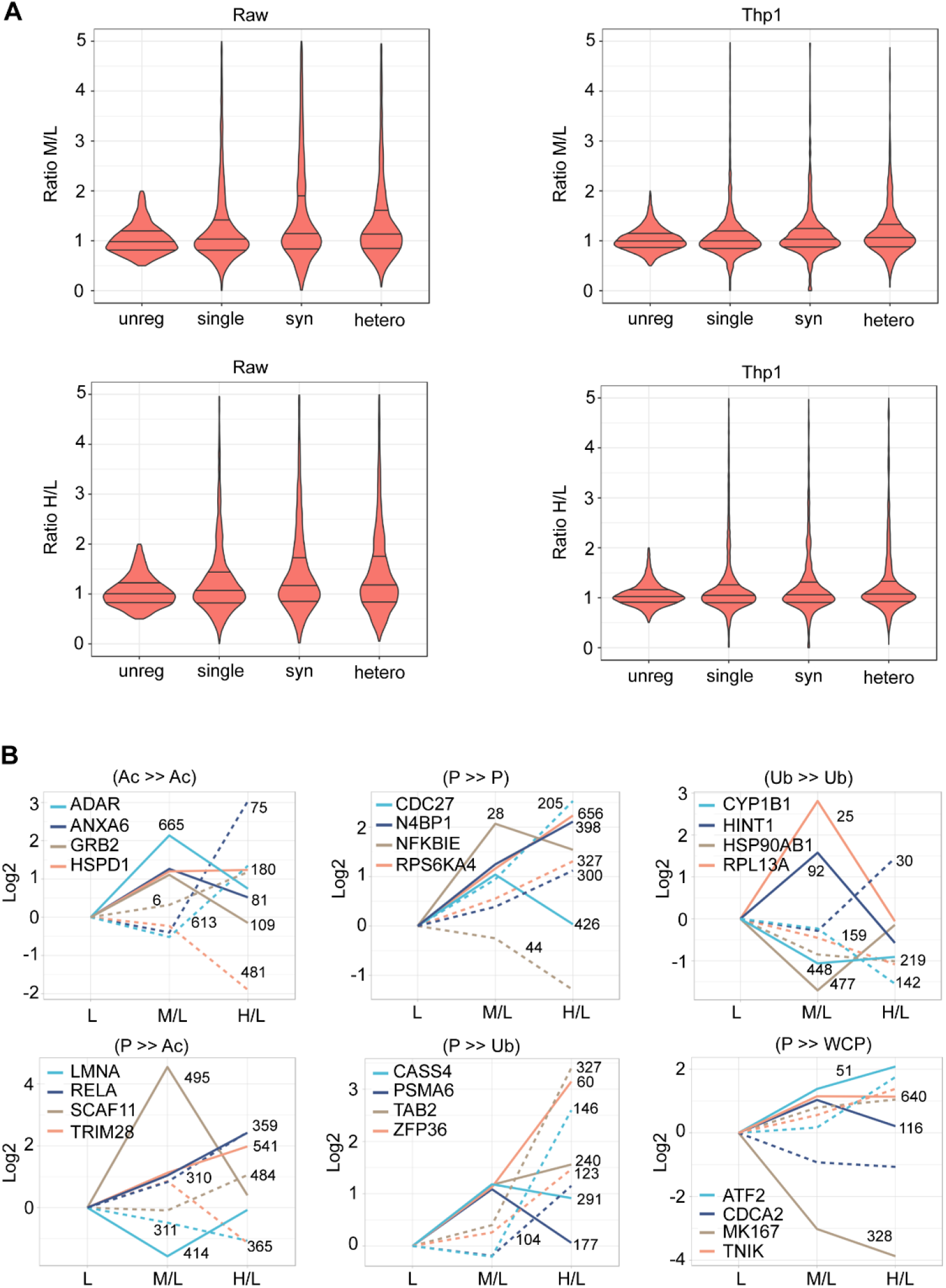
PTM crosstalk between multiple sites on the same protein. (**A**) Distribution of the ratio of all PTM sites identified in proteins in the unreg, single, syn and hetero groups. The lower, median and upper lines in each violin plot correspond to 25%, 50% and 75%, respectively. (**B**) Selected regulated proteins in Thp1 cells belonging to the corresponding category listed on top of each panel. The solid line represents a ‘fast’ regulated event and the dotted line represents a ‘slow’ regulated event. The number next to the line represents the site of PTM on the corresponding protein. The PTM in the left of “≫” is ‘slow’ PTM and The PTM in the right of “≫” is ‘fast’ PTM.

**Figure S6.**
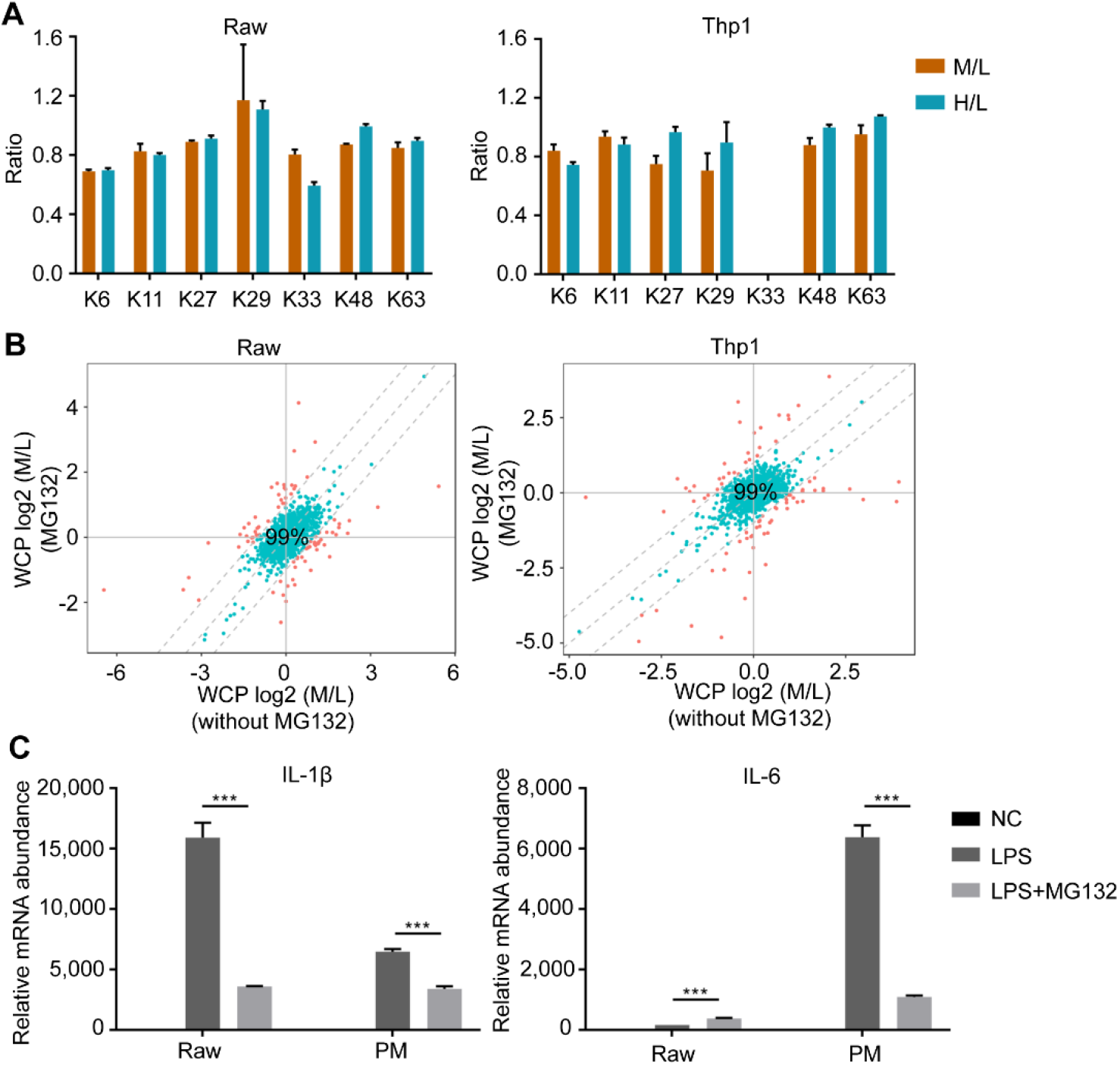
Integrative proteomics reveals a prevalence of both degradative and non-degradative ubiquitylation. (**A**) The ratio of the abundance of ubiquitin lysine sites quantified in the Ub of Raw and Thp1 cells following LPS stimulation with the presence of MG132. (**B**) Comparison of Log_2_ (M/L) values of proteins abundance in the WCP of LPS-stimulated cells treated with or without MG132 for 2 hours. Proteins that exhibited a ≥ 1 Log_2_ (M/L) difference in untreated and MG132-treated cells were considered dramatically affected by MG132 (carmine). (**C**) The relative mRNA levels of *IL-1β* and *IL-6* were generated from the comparison of a certain group with the “NC, negative control”. One representative containing three technical replicates out of two independent experiments is shown. MG132 (2 μM) and LPS were added at the same time. Error bars represent the standard error of the mean and statistical significance was determined by t-test (\**P* ≤ 0.05; \*\**P* ≤ 0.01; \*\*\**P* ≤ 0.001).

**Figure S7.**
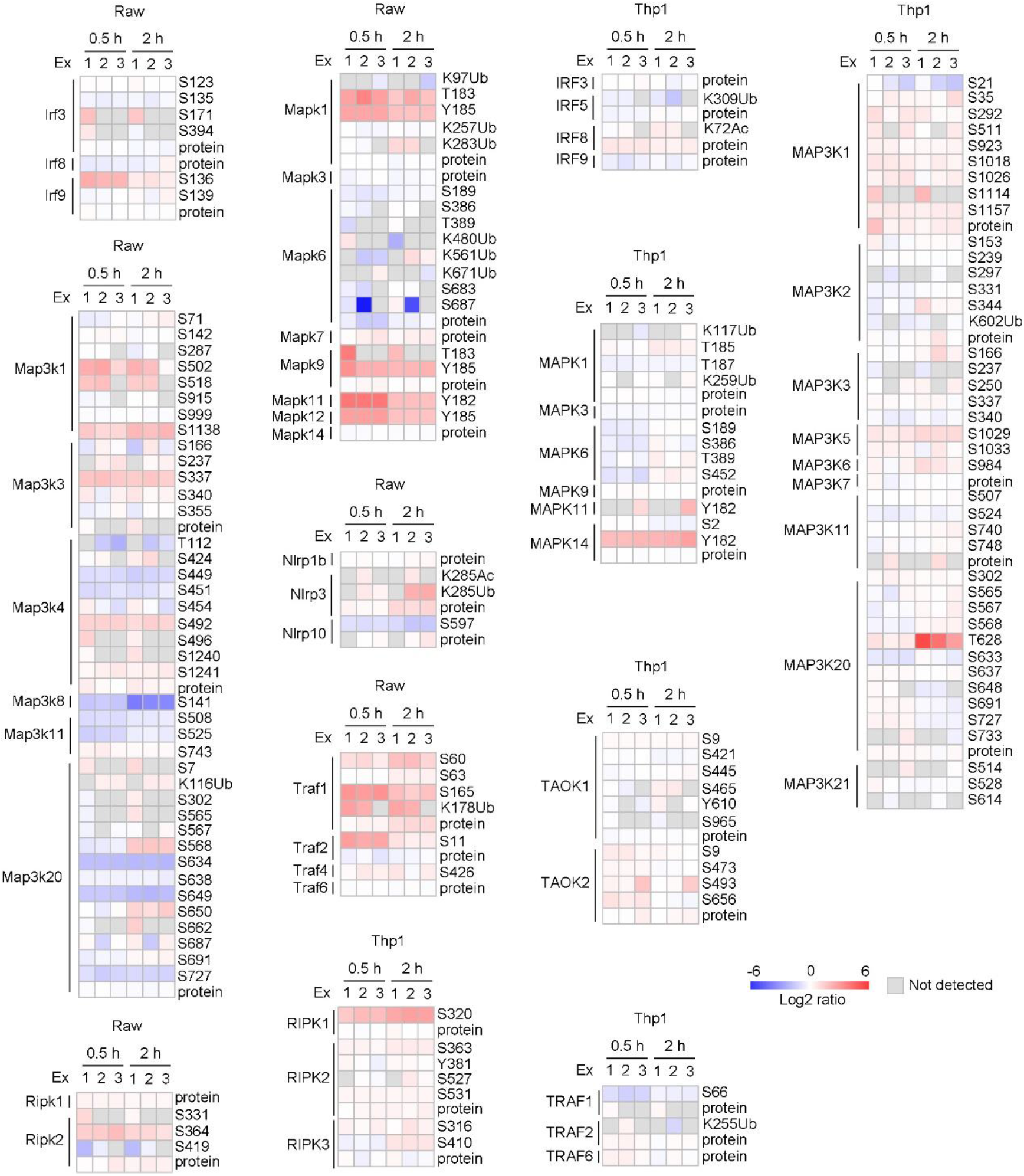
Regulated proteins involved in inflammatory signalling pathways after cells were stimulated with LPS. The diagram shows the intensity of signals for proteins and PTM sites involved in inflammatory signalling pathways identified in cells stimulated with LPS for 0.5 hours and 2 hours.

**Table S1 WCP and PTM data of Raw and Thp1 cells**

**Table S2 Interacting proteins containing PTM crosstalk across proteins**

**Table S3 Time series analysis of PTM data**

**Table S4 siRNA screening results and siRNA sequences**

**Table S5 Conserved proteins between Raw and Thp1 cells**

